# Making yogurt with the ant holobiont uncovers bacteria, acids, and enzymes for food fermentation

**DOI:** 10.1101/2024.09.16.613207

**Authors:** Veronica M. Sinotte, Verónica Ramos-Viana, Diego Prado Vásquez, Sevgi Mutlu Sirakova, Nabila Rodríguez Valerón, Ana Cuesta-Maté, Shannara K. Taylor Parkins, Julia Giecko, Esther Merino Velasco, David Zilber, Rasmus Munk, Sandra B. Andersen, Robert R. Dunn, Leonie J. Jahn

## Abstract

Milk fermentation has a rich history in which food culture, the environment, and microbes intersect. However, the biocultural origins of fermentation practices and microbes have largely been replaced by industrial processes. Here, we consider a historical fermentation originating from Turkey and Bulgaria – ant yogurt. We revisit the traditional practices and modern gastronomic applications that use red wood ants (*Formica rufa* group) to initiate milk fermentation. Subsequently, we characterize the ants and experimental ant-derived yogurts. We uncover that the ant holobiont, which consists of the ants and their microbes, contributes key acids and enzymes to fermentation. Metabarcoding and culturing revealed that lactic and acetic acid bacteria, including species related to conventional sourdough, originate from the live ants and proliferate in the milk. The ants and bacteria consequently introduce formic, lactic, and acetic acid, advantageous for yogurt acidification and coagulation. Last, proteases with the potential to act on casein may alter yogurt texture and are produced by the ants and bacteria. The ant holobiont thus catalyses fermentation akin to the microbial consortia in other ferments. Our findings highlight the value of integrating traditional, gastronomic, and biological frameworks to uncover the origins and applications of microbes for fermented foods.

## Introduction

The fermentation of milk into products such as yogurt, cheese, and kefir originates from ancient practices and has dramatically shaped food cultures. The oldest archaeological evidence for dairying dates to 9,000 years ago in Anatolia (modern-day Turkey) (Evershed et al., 2008). Prehistoric dairy fermentation potentially occurred as early as 7,000 years ago, based on fat and protein residues isolated from ceramics resembling cheese strainers (Evans et al., 2023; Salque et al., 2013). In the millennia to follow, diverse dairy practices transformed milk into a preservable, widespread, and nutritious resource (Fisberg & Machado, 2015; Scott et al., 2022; Warinner et al., 2014). Following suit, dairy fermentation became indispensable to regional cuisines and languages (Multu Sirakova, 2023; Öney Tan, 2010). Yogurt, a tangy fermented milk product, was thus a functional cultural adaptation dependent upon interactions between people, dairy animals, the environment, and most importantly, microbes. It is microbes that enter the milk, and through their enzymatic processes, catalyse the fermentation to acidic, viscous yogurt (Nagaoka, 2019). These interspecies relationships are reflected in the Turkish word for a fermentation starter, *maya*, that ultimately “comes from relations within the broader web of life”, including microbes, animals, plants, and human culture (Multu Sirakova, 2023).

In the early 1900s, microbiologists characterized the first yogurt culture, laying the foundation for a pivotal shift from the diversity inherent in traditional yogurt to a simplified industrialized yogurt. Stamen Grigorov and Ilya Metchnikoff isolated and popularized a species of bacteria from Bulgarian *maya*, *Lactobacillus delbrueckii* subsp. *bulgaricus* (Grigoroff, 1905; Metchnikoff, 1908). Following Metchnikoff, the industrialization of yogurt focused on a small number of bacterial taxa, predominantly *L. delbrueckii* subsp. *bulgaricus* and *Streptococcus thermophilus* (FAO & WHO, 2018). Both species are lactic acid bacteria, which play an important role in industrial and wild fermentations through food preservation, flavour generation, and potential health benefits (Hu et al., 2022; Wang et al., 2021; Zapasnik et al., 2022). However, the focus on a few bacterial species overlooks the biodiversity embodied in traditional yogurts, which can include multiple species and strains (Demirci et al., 2022; Velikova et al., 2018). Revisiting biocultural origins of yogurt fermentation offers opportunities to explore this biodiversity for novel starter cultures (Sedó Melina et al., 2022), to understand the interconnectedness of food systems, and to illuminate the history of yogurt and its cultural praxis.

The bacteria in yogurt stem from the multispecies dimensions of *maya*, which we aim to elucidate for a traditional Turkish and Bulgarian yogurt-making practice - ant yogurt. Generally, the bacterial community that first colonizes the milk and establishes the fermentation ecosystem may stem from the dairy animal (Wouters et al., 2002), the person making the ferment (Kothe et al., 2024; Reese et al., 2020), or the environment, such as vegetation (Michaylova et al., 2007), air, or containers used. Yogurt cultures are then propagated by adding a small amount from a primary or old yogurt to new milk, a process referred to as “backslopping.” A historic practice for starting the first or primary yogurt in Turkey (Öney Tan, 2010) has been documented by the ethnographer Ali Rıza Yalman, “If the nomads want to make yogurt and cannot find enough starter culture to make yogurt, they crush the tiny eggs of the ants sheltering under the stones in their palms. When you put this into the milk […], that milk becomes yogurt.” (Yalman, 1977). In neighbouring Bulgaria, a traditional spring practice involves fermenting yogurt within a red wood ant colony (Multu Sirakova, 2023). Further oral histories from Sharri mountains of Albania and North Macedonia also recall the use of ants in traditional yogurt fermentation (F. Demiraj, personal communication, May 2023). These countries are connected by cultural threads preceding the establishment of national boundaries, contributing to general continuity between their culinary practices, both today and across millennia. The ethnographic evidence of ant yogurt across regions suggests ants may play an overlooked functional role in fermentation.

Here, we test the hypothesis that yogurt fermentation can be initiated by the ant “holobiont”, which includes both the ant and the microbial communities inherent to it (Bordenstein & Theis, 2015; Margulis & Fester, 1991). The ant holobiont may contribute acids and enzymes key to the fermentation (Figure 1). First, we explored the potential of red wood ants (*Formica rufa* group) (Robinson & Stockan, 2020) as starter cultures for yogurt based on ethnographic accounts and as culinary ingredients for modern gastronomy. Then, we characterized the microbial community of the ants *F. rufa* and *F. polyctena* and yogurts derived from *F. polyctena*, quantified organic acids in the yogurts, and assessed the proteases and peptidases originating from ants and bacteria (Figure S1). The study overall examines an overlooked ecological niche of bacteria in the context of traditional practices to identify microbes key to past and future fermented foods.

**Figure 1.**
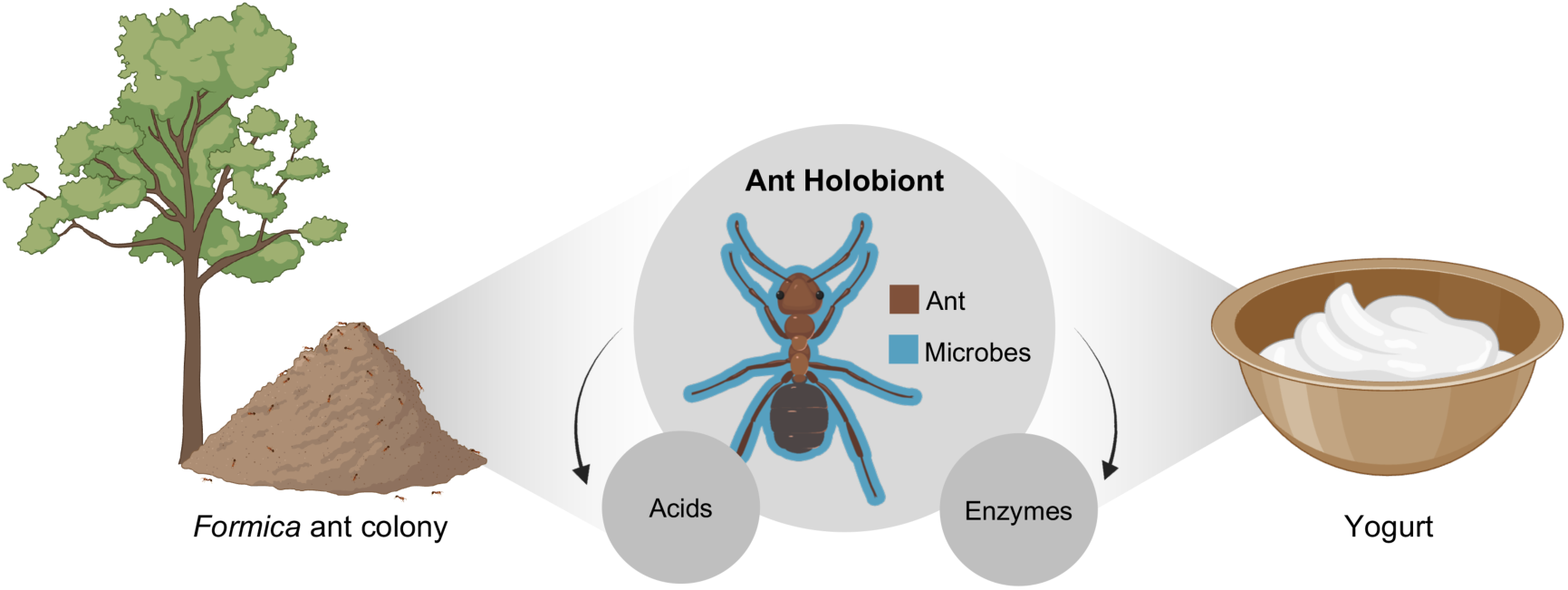
Hypothesized contributions of the ant holobiont to yogurt fermentation. *Formica* red wood ants create characteristic thatched mounds (left), which may have been used in traditional yogurt fermentation practices (Mutlu Sirakova 2023). The traits of the ant holobiont, the composite of both ant and the microbes found within it, are hypothesized to act as the starter of the yogurt fermentation (centre). These include the microbes (lactic acid and acetic acid bacteria) that are found in and on the ants, acids that are produced by the ant and the ant-associated microbes, and enzymes that have ant or microbial origins.

## Results

### The ant holobiont serves as a catalysing agent for dairy applications

We worked towards a holistic understanding of traditional uses of ants as a yogurt starter by conducting fieldwork in Bulgaria in a community that retained an oral history of the practice (Multu Sirakova, 2023; Sinotte et al., 2023). This community also is the ancestral home of one of the authors (S. Mutlu Sirakova). We used an ant colony selected by members of the community, which was the species *F. rufa*. This is one of four species in the *F. rufa* group that maintain species ranges within Bulgaria (Antonova & Marinov, 2021). To make the yogurt, four live ants were added to the jar of warmed raw milk (Figure 2A), a cheese cloth was placed over the top of the glass jar, and the jar was left to ferment within the colony overnight (Figure 2B). Here, the ant could act as an inoculum, and the colony could serve as an incubator since the nest itself is known to produce heat (Coenen-Stass et al., 1980). The jar was retrieved after a day of incubation. The acidity, texture, and flavour were indicative of early stages of yogurt fermentation. We observed that the milk had acidified to pH 5, coagulated at the bottom of the container (Figure 2C), and “had a slight tangy taste with mild herbaceousness and pronounced flavours of grass-fed fat” (D. Zilber).

**Figure 2.**
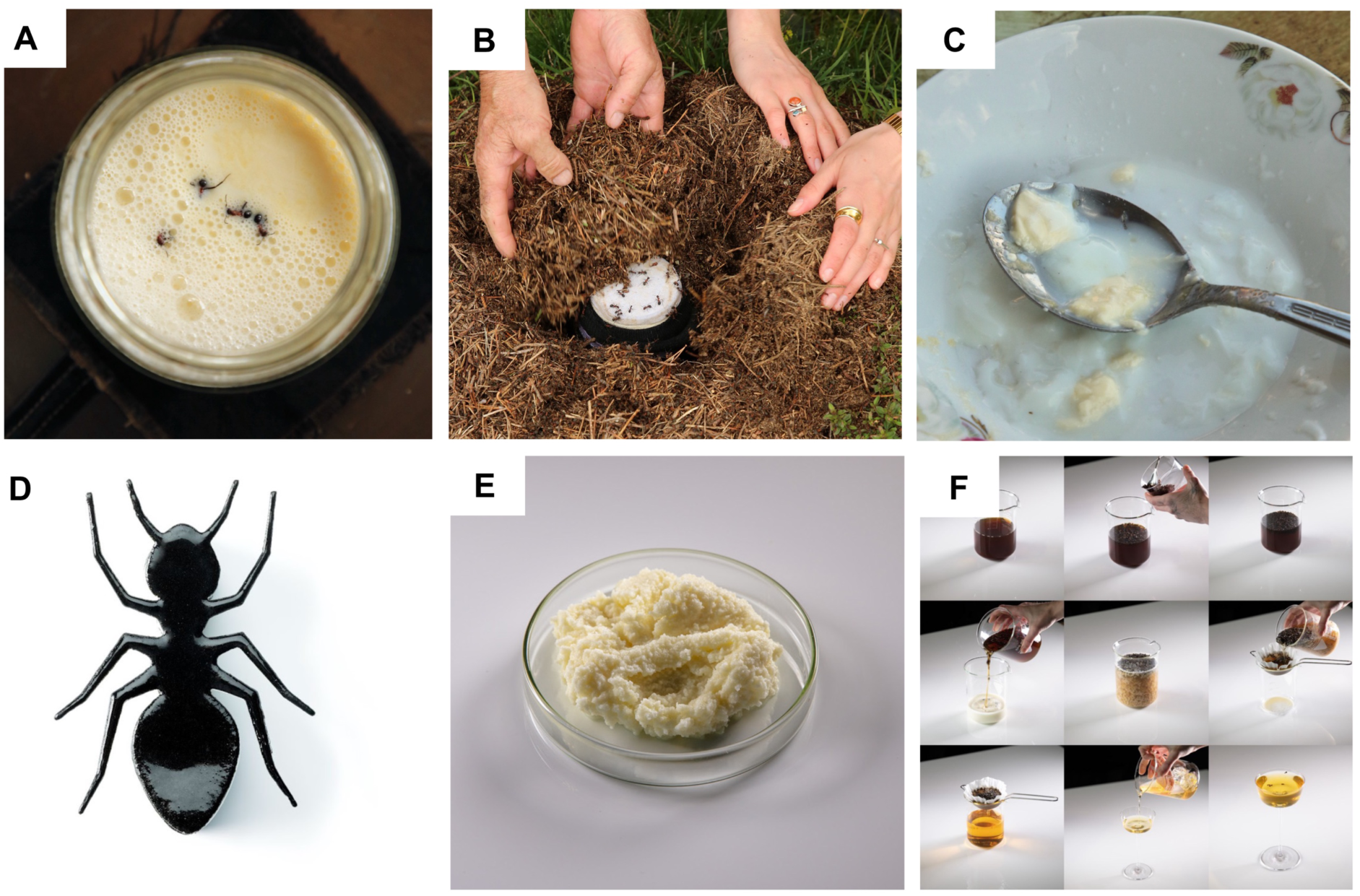
The ant holobiont serves as a catalysing agent for dairy applications. (**A-C**) Photographs of traditional yogurt fermentation initiated by *Formica* ants taken during field work in Bulgaria. (**A**) Ants were added to warmed milk that was then (**B**) buried within the ant colony and left to ferment overnight. (**C**) The resulting milk had started to coagulate and acidify, indicative of early stages of yogurt fermentation. (**D-F**) Culinary applications created by restaurant Alchemist’s research and development team using *F. rufa* ants: (**D**) ant yogurt ice cream sandwich, (**E**) ant “mascarpone-like” cheese, and (**F**) milk-wash cocktail (Photos a-c by David Zilber, and d-f by Søren Gammelmark and Kåre Knudsen of Restaurant Alchemist).

Culinary applications of the *F. rufa* ants were created by the research and development team at Restaurant Alchemist, ranked number eight in the world (50 Best, 2024), holding two Michelin stars (Guide, 2024), and known for science-centred innovation. Co-authors based these applications on previous experiments where the addition of ants to milk caused coagulation. Presumably, this coagulation was due to the *F. rufa* ants’ formic acid. Discussions on the traditional uses of ants to coagulate milk (T. Tan, personal communication, April 2022) further directed the development of culinary applications. Thus, the innovations leverage the potential of ant-derived formic acid and microbe-derived organic acids to coagulate milk.

Three culinary applications were developed using live ants and ants that were frozen and subsequently dehydrated. First, the “ant-wich” contained ice cream derived from sheep yogurt made with live ants as starter culture (Figure 2D). The ants provided a distinct, pungent acidity that contrasts with the fat of the milk, and a serving temperature of -11°C balances the desert. Second, a goat milk “mascarpone” was developed using dehydrated ants to catalyse the milk coagulation (Figure 2E). The texture was like commercial mascarpone, yet the flavour was pungent and aromatic, like a mature pecorino cheese. Third, a milk wash cocktail was created, which is a dairy-based cocktail dating to the early 1700s (Wondrich, 2010). Classically, the milk is curdled with acid from citrus and subsequently filtered to remove the dairy solids, resulting in a clear beverage with more richness and body. In this case, dehydrated ants induced the curdling and separation of milk (Figure 2F). The cocktail had fruity notes from the apricot liqueur and the brandy (raisins, dried figs, and caramelized apples), and a silky texture from the residual milk whey. Replacing citrus with ants resulted in a milder acidity and a distinct flavour with additional fruity notes. Despite the potential of these culinary innovations, we note that ants are not included among the four insects authorized for sale as a food product in the European Union according to Regulation 2015/2283 on Novel Foods.

Given that ants have diverse culinary applications with potential to initiate fermentation, we aimed to further elucidate the role of the ant holobiont. We hypothesized that the ant holobiont, consisting of the ant and its microbial partners, contributes to fermentation with bacteria, acids, and enzymes. To address this hypothesis, we created three elaborations of fermented milk with live, frozen, or dehydrated ants, per the modern gastronomic applications. The ants were collected in late spring and early autumn to further determine the impact of the season on the microbiome and the fermentation.

### Live ants provide small, stable, and controlled yogurt microbiomes

We first characterized the bacterial microbiome of red wood ant sister species *F. rufa* and *F. polyctena* (Borowiec et al., 2021) with 16S *rRNA* metabarcoding. Our aim was to examine how the ant yogurt practice may be translated across species and geographies. These ants have ranges spanning Denmark, where experimental yogurts were made, and Bulgaria or Turkey (Guénard et al., 2017; Janicki et al., 2016), where ethnographic histories originated (Figure 3A). Both species are found along the edges of pine forests, have thatched pine needle mounds, and near-identical morphological characteristics (Seifert, 2021). These similarities make them all but indistinguishable in the field to ant experts (Seifert, 2021), suggesting both may have been used in traditional and modern culinary applications.

**Figure 3.**
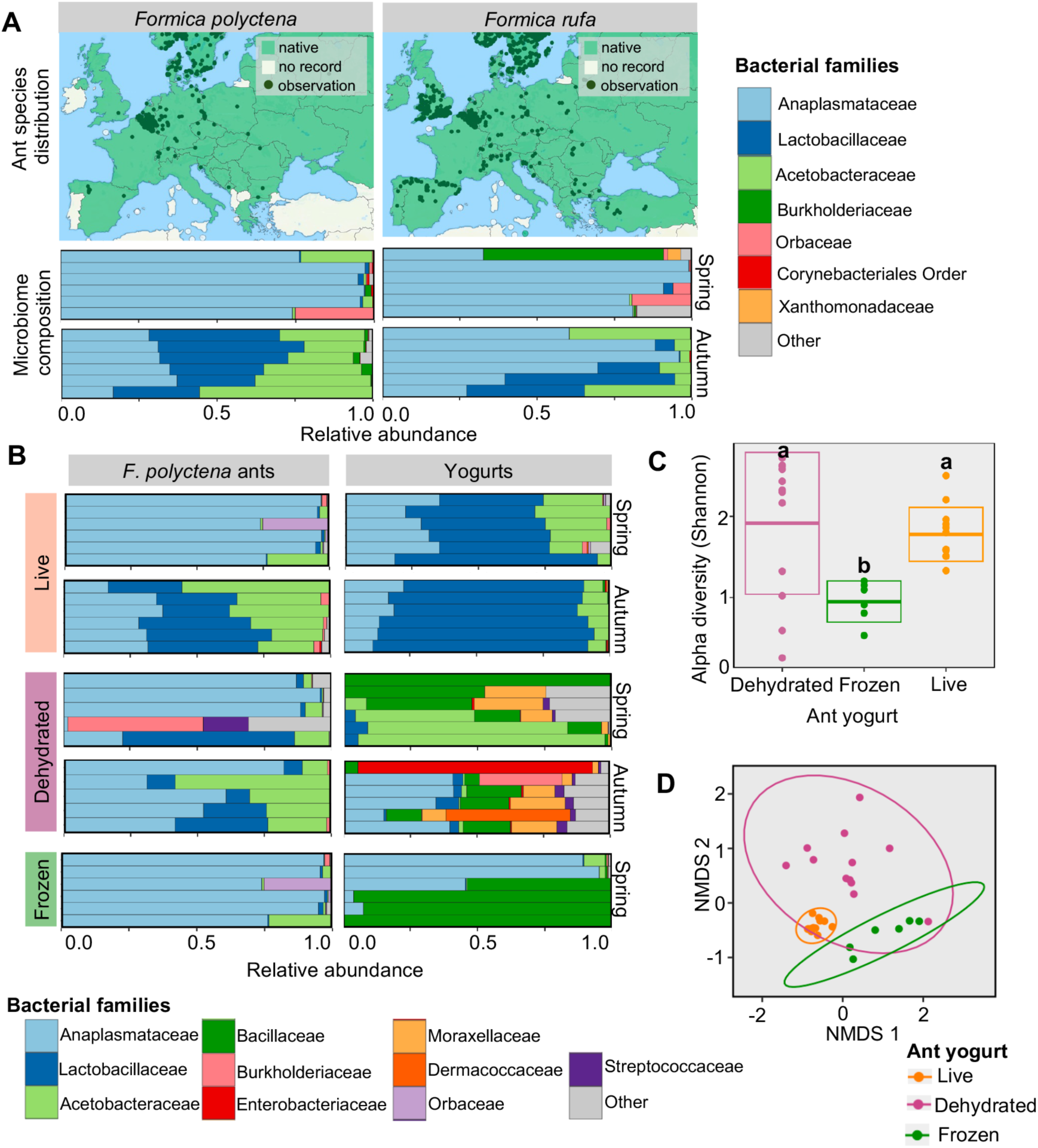
Live ants provide small, stable, and controlled yogurt microbiomes. (**A**) European species ranges of *F. polyctena* and *F. rufa* (top), both within the *F. rufa* group of red wood ants (maps modified from antmaps.org). The microbiome composition of ant species across seasons (bottom). Bar represents four pooled ants from a colony of each respective species, collected in Denmark. The top 8 bacterial families among the samples are illustrated. (**B**) The bacterial microbiome of the three ant preparations and corresponding ant yogurts made in spring and autumn. Bars represent replicates, and the top ten bacterial families are shown. Live ant microbiomes are also represented in panel (**A**), and live and frozen ants are identical, as they live ants that were frozen prior to extraction are also representative of the frozen ants themselves. They are shown for direct comparison to the corresponding yogurts. (**C**) The alpha diversity of ant yogurts. Letters indicate pairwise statistical differences (Tukey HSD: *p*<0.05), and boxes illustrate standard deviations. (**D**) The beta diversity of ant yogurts based on Bray-Curtis distances and Non-Metric Multidimensional Scaling (NMDS) analysis. The grouping is supported by a PERMANOVA.

The microbiomes of the two *Formica* species were dominated by lactic acid bacteria (Lactobacillaceae), acetic acid bacteria (Acetobacteraceae), and obligate intracellular bacteria (Anaplasmataceae) (Figure 3A). The bacterial families and genera (Figure S2A) align with *Formica*-specific microbiomes associated with species across the Northern Hemisphere (Jackson et al., 2023; Jackson et al., 2022; Kaczmarczyk-Ziemba et al., 2020; Zheng et al., 2022). *Formica* ant microbiomes were seasonal. We observed an increase in relative abundance of lactic and acetic acid bacteria (Figure 3A) and overall microbial biomass (i.e. load) in the autumn, compared to spring (Figure S2B). Seasonal microbiomes have not been previously documented in *Formica* ants and are germane to the use of these ants in fermentation. Overall, the consistent presence of lactic and acetic acid bacteria in the *F. rufa* group indicates the ant microbiome may be pertinent to food fermentation.

We then tested the hypothesis that lactic and acetic acid bacteria in the ants may transfer to the yogurt. Based on 16S *rRNA* metabarcoding of experimental yogurts made with *F. polyctena* under aseptic conditions, we confirmed our hypothesis that bacteria from the ants contribute to the yogurt microbiome. The preparation of the ants, namely whether they were live, dehydrated, or frozen, distinctly impacted the bacteria communities in the resulting yogurts (Figure 3B). Like the ants themselves, the live ant yogurts were consistently dominated by the lactic and acetic acid bacteria, which displayed seasonal differences in composition. Here, the most abundant bacterial genus was *Fructilactobacillus* (Figure S2C). We note that these true abundances may have been higher, considering our metabarcoding positive control indicates a slight underestimation of the abundance of lactic acid bacteria in samples (Figure S3). In contrast, dehydrated ant yogurts exhibited variable microbiomes. Frozen ant yogurts contained two bacteria groups: Bacillaceae, which can proliferate in the yogurt, and Anaplasmataceae, which obligately lives in ant cells (Buchner, 1965; Jackson et al., 2022) and thus cannot grow in the yogurt. Here, dehydration and freezing likely reduced bacterial viability (Santivarangkna et al., 2011) and consequently favoured stochastic community assembly or freeze-resistant Bacillaceae (Nicholson et al., 2000). The persistence of lactic and acetic acid bacteria in live ant yogurt suggests it is the best starter for fermentation and provides biological support for the traditional, live ant-based yogurts.

We determined the consistency of the bacterial microbiome composition across the yogurts. Alpha and beta diversity metrics were used, where alpha diversity is the number of bacterial strains per sample (i.e. amplicon sequence variants) and beta diversity is the difference in the composition of strains from one sample to the next, while weighting relative abundance. The ant preparations significantly affected the alpha and beta diversity of the yogurts (LM: F_2,27_=5.387, *p*=0.0108, based on Shannon index; Adonis: F_2,27_= 8.4402, *p*<0.0001, based on Bray Curtis distances). The alpha diversity of the live ant yogurts was intermediate (Figure 3C), but the beta diversity was very low (Figure 3D). Therefore, the live ant yogurts tended to have the same species and composition from one sample/preparation to the next, as one would hope to see in predictable and controlled fermentations.

### Bacteria from the ants proliferate in the milk

We hypothesized that bacteria from the ant holobiont grew in the milk, facilitating fermentation. To assess growth, we determined the bacterial load in ants and yogurts with quantitative PCR (qPCR) and identified viable bacteria with culturomics using 16S *rRNA* Sanger sequencing. Live ants introduced lactic acid bacteria that proliferated in the yogurt. In the spring, yogurts contained a higher loads of lactic acid bacteria than the ants introduced as a starter (Figure 4A; *t* (2, 10) = - 2.975, *p*=0.0139). However, in autumn, we observed the opposite pattern (Figure 4A; *t* (2, 10) = 3.269, *p*=0.0084), suggesting that not all lactic acid bacteria from the ants enter the milk, or that the bacteria associated with the autumn ants are less prolific. While lactic acid bacteria dominated live ant yogurts, it made up a marginal amount of the bacterial load in frozen and dehydrate yogurts (Figure S4A).

**Figure 4.**
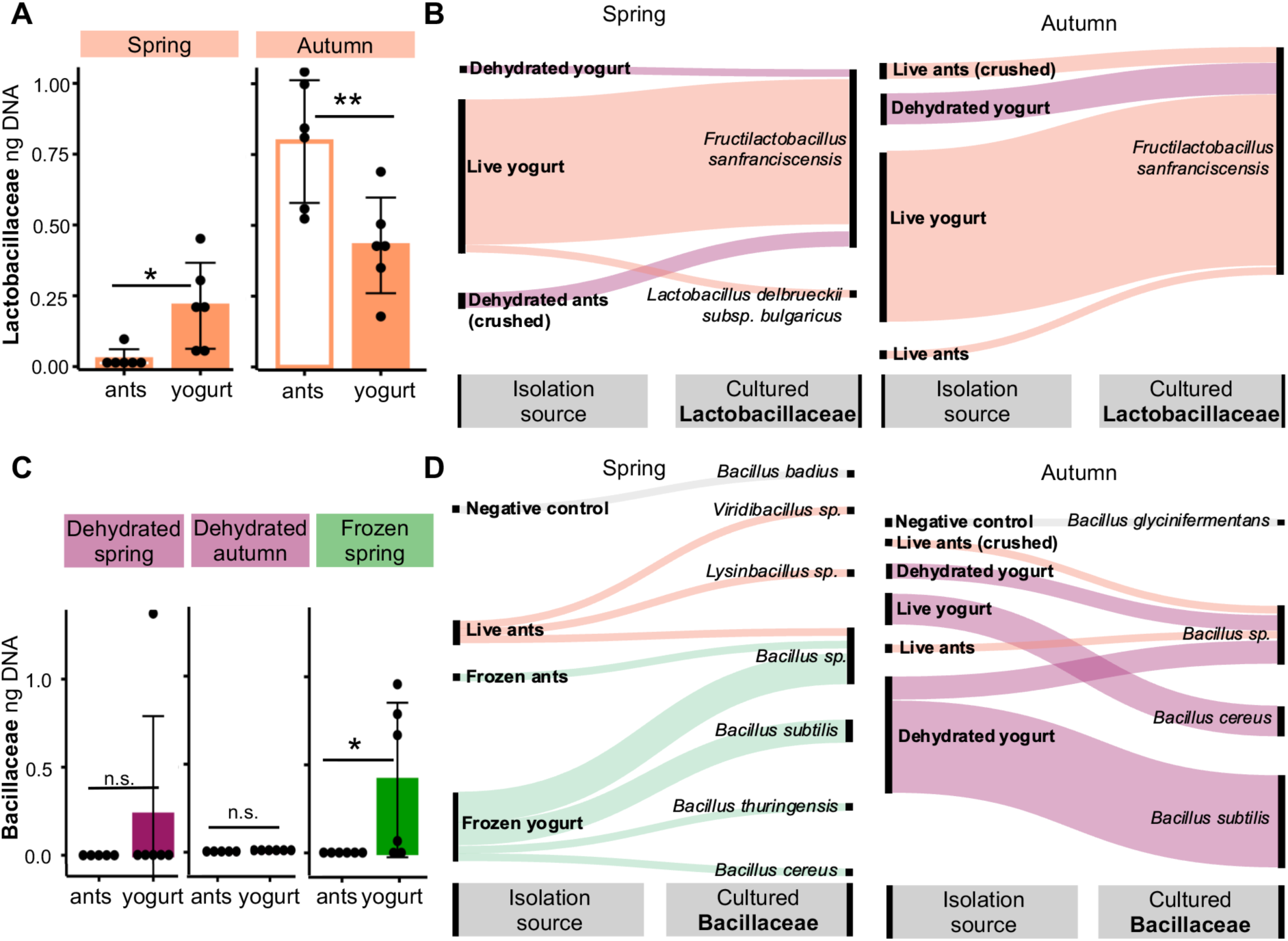
**Bacteria from the ants proliferate in the milk**. (**A**) The estimated amount of lactic acid bacteria (Lactobacillaceae) DNA, representative of lactic acid bacterial load, in the live ants and resulting yogurts across seasons. (**B**) The diversity of culturable species of lactic acid bacteria from the ants and yogurts, where the majority of isolates stem from live ants and live ant yogurts. The thickness of each line approximates abundance of culturable bacteria by representing the number of microbial media plates where the bacteria grew compared to other samples. (**C**) The amount of DNA from spore-forming Bacillaceae, indicative of Bacillaceae bacterial load, is exceptionally low in dehydrated and frozen ants and increases in the respective yogurt fermentations. (**D**) Culturable species of Bacillaceae isolated from ants, yogurts, and negative control yogurts. Frozen ant yogurts were not made in autumn. Line thickness represents an approximation of culturable bacteria as above. (* = *p* ≤ .05; ** = *p* ≤ .01; *** = *p* ≤ .001; n.s. = p>0.05; y-axis present on a logarithmic scale).

Yogurt made with live ants contained several species of culturable lactic acid bacteria (Figure 4B). Aligned with the dominance of *Fructilactobacillus* in the community analysis, we isolated *F. sanfranciscensis*, a bacterium not only associated with ants (Zheng et al., 2022) but also sourdough bread fermentation (Gänzle et al., 2023; Kline & Sugihara, 1971). We further cultured and isolate of *Lactobacillus delbrueckii* subsp*. bulgaricus*, typically found in fermented dairy products (FAO & WHO, 2018). Last, we isolated the acetic acid bacteria, *Oecophyllibacter saccharovorans* (Figure S5), previously characterised in association with ant genera closely related to *Formica* (Blaimer et al., 2015; Brown & Wernegreen, 2016; Chua et al., 2018). The diversity of culturable lactic acid bacteria thus indicates a niche overlap between ants and fermented foods.

In the yogurts made with dehydrated and frozen ants, the bacterial load disproportionately came from spore-forming Bacillaceae. Notably, there is little to no Bacillaceae biomass in the ants, especially when compared to the yogurts (Figure 4C; *t*_frozen_ (2, 10) = -2.307, *p*=0.0437), and this Bacillaceae did not proliferate in live ant yogurts (Figure S4B). We cultured several species of Bacillaceae, stemming largely from dehydrated or frozen ants or yogurts (Figure 4D). This included the food contaminant, *Bacillus cereus* (Dietrich et al., 2021). Although we cannot determine if the *Bacillus* load was sufficient to be problematic for consumption, it still indicates a potential risk. Therefore, we conclude that the microbiome of dehydrated and frozen ants and their corresponding yogurts are undesirable for food fermentations.

### Ants and bacteria contribute to the acidification of yogurt

We hypothesized that the ant holobiont, composed of both the ant and its microbes, may contribute acids key to fermentation. In yogurt, bacteria metabolize lactose and consequently produce high amounts of lactic acid and often low amounts of acetic and formic acid (Sieuwerts et al., 2010). These acids are essential to the tangy flavour, thick texture, and preservation of yogurt (Fernandez-Garcia & McGregor, 1994). Moreover, *Formica* ants (though, notably, not all ants) have a venom gland that contains largely formic acid that may be up to 10% its body weight (O’Rourke, 1950), although the whole ant contains a more complex mélange of chemicals (Löfqvist, 1976). Thus, the acid from the ant holobiont may have three effects. First, it may engender yogurt tastes and textures, even without the effects of microbes. Second, it can create acidic conditions that favour acid-producing microbes. Third, the acid-producing microbes, independent of the ant, can themselves alter the acidity and flavour of the ferment. To investigate if the components of the ant holobiont, body and microbes, contribute organic acids to the yogurt, we quantified formic, lactic, and acetic acid in yogurts and controls by HPLC and measured pH before and after fermentation (Figure 5).

**Figure 5.**
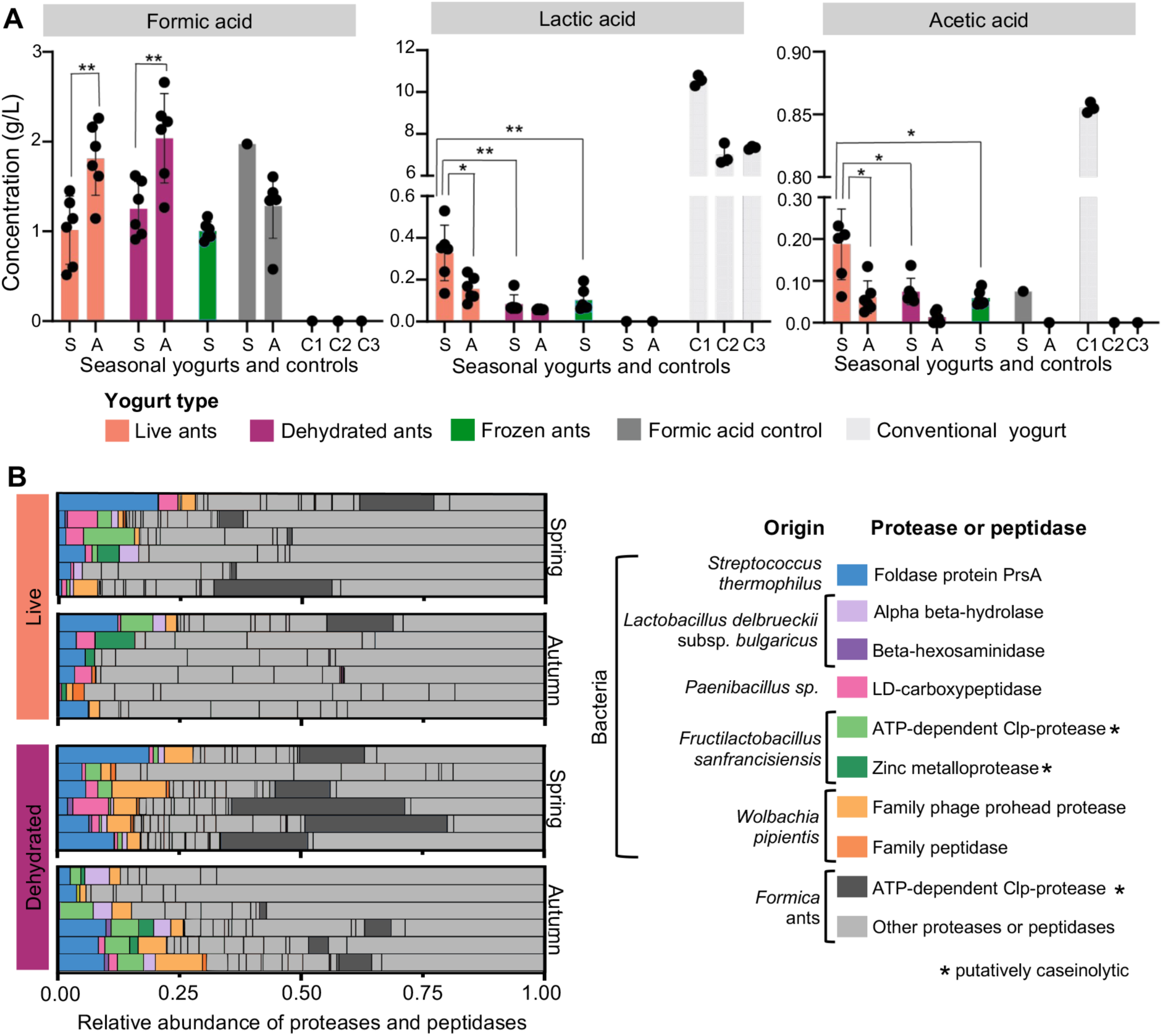
Organic acids, proteases, and peptidases in ant yogurts. (**A**) Concentration of metabolites formic acid, lactic acid, and acetic acid in the yogurts from spring (S) and autumn (A). The samples include yogurts made with live, dehydrated, and frozen ants, formic acid, and three conventional yogurts as controls (C1-3). Data points represent biological replicates and error bars represent the standard deviation. Statistical analysis was performed using Student’s t-test (two-sample unequal variance; * = P ≤ .05; ** = P ≤ .01; *** = P ≤ .001). (**B**) Proteases and peptidases detected in live and dehydrated ant yogurt samples. Proteases and peptides originate from the ant holobiont, including the ant and its bacteria. It also includes bacteria found in conventional yogurt, that were potentially present in low abundances. The rows represent biological replicates. Milk proteins were removed to improve the visualization of the proteases and peptidases. Milk proteins made up 96.19 ± 1.07% of the samples. The absolute abundance of all proteins in the proteomes can be found Table S2.

The ant holobiont contributed formic acid, which was the most abundant acid found in the yogurts (Figure 5A). The amount of formic acid was in a similar range to that in the control, in which formic acid alone was added to pH 4.6 to induce coagulation. The season, but not the type of yogurt (live, dehydrated, frozen) further dictated the formic acid levels (LM results). The higher levels in autumn (t-test: *t*_live_ (2, 10) = -3.500, *p*= 0.0057; *t*_dehydrated_ (2, 10) =-3.297, p=0.0081) may be due to the seasonality of formic acid production (O’Rourke 1950). Therefore, the acid from the ants entered the milk and likely shaped the unique acidic flavour and texture of the culinary applications, contrasting the rounded flavour of lactic acid typically predominant in fermented dairy.

Lactic and acetic acid were also detected in the ant yogurts, validating that ant-associated bacteria likely contributed acids during fermentation (Figure 5A). Acid profiles align with the microbiome data. Lactic and acetic acid bacteria were the most abundant genera in live ant yogurts (Figure 3A). Similarly, these acids are highest in live ant yogurts compared to dehydrated (*t*_lactic_ (2, 10) =4.270, *p*=0.0052; *t*_acetic_ (2, 10) =3.065, *p*=0.0220) and frozen yogurts (*t*_lactic_ (2, 10) =3.842, *p*=0.0060; *t*_acetic_ (2, 10) =3.653, *p*=0.0147). In live yogurts, we observed lactic and acetic acid were higher in spring (*t*_lactic_ (2, 10) =2.897, *p*=0.0231; *t*_acetic_ (2, 10) =3.325, *p*=0.012), the season in which we isolated the greatest diversity of lactic acid bacteria (Figure 4C). This suggests different strains of bacteria may inherently vary in acid production. Notably, ant yogurts had less lactic acid, acetic acid (Figure 5A), and bacterial biomass (Figure S6A) than conventional or homemade yogurts. We did not establish and perform the optimal fermentation conditions for the ant yogurt, and thus the fermentation was incomplete. However, we can conclude that the ant microbes contribute organic acids to fermentation.

Ant yogurts contained on average 2.5 g/L total organic acids, while conventional yogurts contained up to 12 g/L total organic acid. Thus, the ant yogurt remained in a pH range of 5.0-5.9 and did not reduce to a pH of 4.2 of conventional yogurt (Figure S6B), given the conditions and timing of our experiment. Initially, the pH of the milk dropped upon the addition of ants, likely due to the formic acid from the ants (Table S1). The pH continued to decrease through the fermentation (Figure S6B), likely because of acid-producing microbes. Typically, in yogurt fermentation, a drop to pH 5.3 initiates coagulation, and a further acidification to pH 4.6 completes coagulation by causing aggregation of casein micelles (Robinson et al., 2006) The ant yogurt coagulated yet largely did not reach such a low pH. This suggests another aspect of the fermentation denatured caseins, namely proteases.

### Ants and microbes contribute proteases and peptidases, potentially modifying yogurt texture

Our final hypothesis was that the ant holobiont may contribute enzymes, in particular proteases, that modify the yogurt texture. Proteases can differentially cleave and degrade casein, the main protein in milk, and result in a firm or fluid yogurt texture (Amani et al., 2017; Gassem & Frank, 1991). Based on the pH and coagulation results, we predicted the ants themselves may contain milk-clotting proteases, like those previously identified in mealworms (Yang et al., 2022). Casein proteases and peptidases are also typically produced by the bacteria in yogurt (Amani et al., 2017). Thus, the ant holobiont, ants and bacteria, may contribute proteases to the yogurt fermentation. To test this hypothesis, we conducted an untargeted proteomics analysis based on a self-curated database that contained proteases from *Formica* ants, ant-yogurt bacteria, conventional yogurt bacteria, and milk proteins.

We confirmed our hypothesis that the ant holobiont contributes proteases and peptidases to the yogurt fermentation (Figure 5B). Several ant proteases were identified in the yogurt (Figure 5B), one of which, an ATP-dependent CLP protease, is known to have caseinolytic potential (Maurizi et al., 1990). Ant-associated bacteria further contributed proteases and peptidases. The lactic acid bacteria *F. sanfranciscensis* produced two proteases, ATP-dependent CLP protease and zinc metalloprotease, both known to have caseinolytic homologues (Jia et al., 2015; Maurizi et al., 1990). Proteases and peptidases typically belonging to bacteria in conventional yogurt, *L. delbruecki*i subsp. *bulgaricus* and *Streptococcus thermophilus*, were found, as well as ant yogurt bacteria *Paenibacillus* and *Wolbachia pipentis*. Last, we identified milk proteins such as casein, which made up a high percentage 96.19 ± 1.07% of the proteome of samples (Table S2), potentially because of the low bacterial load compared to conventional yogurt. There were no significant differences in the relative abundance of all proteins in the proteome between live and dehydrated ant yogurts or seasons (Figure S7). Thus, proteases in both live and dehydrated yogurts may, in part, catalyse coagulation in the experimental yogurts and in the modern culinary applications. Taken together, the ant holobiont, ants and microbes, contribute proteases potentially relevant to food fermentation.

## Discussion

In this study, we explored the traditional practice of using ants as a starter for yogurt fermentation. We hypothesized that the ant holobiont, both the ant and its microbes, catalysed the fermentation. First, we uncovered bacteria from the ants shaped the yogurt microbiome. Using live ants as a starter, in contrast to frozen or dehydrated ants, resulted in a consistent microbiome containing lactic and acetic acid bacteria, desirable for fermentation. These ant-bacteria included some species thought of as canonical fermentation microbes, such as *F. sanfranciscensis*. Second, we discovered that the addition of the ant holobiont to milk introduced acids and proteases. Formic acid originated from the ant, while lactic and acetic acid were likely produced by bacteria. The ants and bacteria also contributed proteases with caseinolytic potential. Our results revealed that the ant holobiont can contribute to fermentation and thus prompted reconsideration of the microbial consortia in our ferments, their origins, and applications to fermented food.

We uncovered how ants and their bacteria synergistically shape the fermentation by applying the holobiont framework. Holobiont theory considers a host and its associated microbes collectively, where the interplay between animal and microbes shapes the resulting ecology and evolution of the whole, host plus microbes (Bordenstein & Theis, 2015; Margulis & Fester, 1991; Zilber-Rosenberg & Rosenberg, 2008). When animals or plants are used in fermentation, their holobionts may structure that fermentation in ways that reach beyond the effect of the individual parts. In this study, it was thus essential to include the entire ant holobiont, rather than isolating the bacteria alone for fermentation. *Formica* ants produce formic acid for defence (O’Rourke, 1950; Tragust et al., 2020) and feed on the sugar-rich honeydew (Ivens & Kronauer, 2022) and protein-rich insects (Robinson & Stockan, 2020). In response, the lactic and acetic acid bacteria are hypothesized to metabolize the honeydew (Zheng et al., 2022), are highly acid tolerant (Tragust et al., 2020) and are potential contributors to proteolysis. Once ants are introduced into milk, the properties of the ant holobiont extend to the yogurt ecosystem. Formic acid initially acidifies the milk, ant-bacteria break down milk sugars producing additional acid, and proteases liberate peptides and amino acids for bacterial growth. By looking through the lens of the holobiont, we could clarify why *Formica* ants and bacteria are pre-adapted for the fermentation of yogurt and potentially other foods.

The effect of the ant holobiont on yogurt fermentation parallels the role of microbes in other fermented foods. In most ferments, multiple species of microbes offer complementary contributions to the fermentation process (Canon et al., 2020; Sieuwerts et al., 2008). In conventional yogurt, the bacteria *S. thermophilus* produces formic acid that is then metabolized by *L. delbrueckii* subsp. *bulgaricus*, and in return, *L. delbrueckii* subsp. *bulgaricus* produces proteases. Collectively, these bacteria then metabolize lactose and produce lactic acid to transform milk into tangy yogurt (Sieuwerts, 2016; Sieuwerts et al., 2010). In sourdough, yeast and lactic acid bacteria can secrete factors that promote one another’s growth (Sieuwerts et al., 2018), and overall have complementary roles in the fermentation of sugars and protein derivatives (Gänzle, 2014; Gänzle et al., 2023). Together, sourdough microbes create leavened bread with acidic, kokumi, and umami flavours (Gänzle et al., 2023). In the ant yogurt, our results suggest the separate elements of the ant holobiont, the ant body and microbes, act synergistically. Consequently, the ant holobiont introduces acids, proteases, and potentially other compounds that contribute to the organoleptic characteristics such as distinct acidity, coagulation, and flavour.

Intriguingly, we isolated a species of lactic acid bacteria from the ant yogurt that is commonly found in fermented foods, *F. sanfranciscensis*. The bacterial species *F. sanfranciscensis,* is one of the most prevalent bacteria in sourdough bread globally (Gänzle & Zheng, 2019; Landis et al., 2021). The *Fructilactobacillus* clade appears to have diversified with ants (Jackson et al., 2023) and related social insects (Duar et al., 2017). We suspect that *F. sanfranciscensis* has evolved in ants over millions of years and was only introduced into fermented foods with the advent of bread making in the last several thousand years (Arranz-Otaegui et al., 2018; Carretero et al., 2017). In the future, the bacteria isolated from these ants may be assayed for key functions in ferments, such as sugar metabolism and exopolysaccharide formation. These results lay the groundwork for unveiling the diversity of fermentative bacteria residing in ants and for elucidating the bacterial evolution from ant holobionts to fermented foods.

Our results bring forth new questions we believe are central to understanding the ecological and ancient origins of fermentative microbes. The ecological origins and assembly of ferment microbiomes, which colonize fermentations in lieu of backslopping, are only beginning to be characterized (Louw et al., 2023). Recently, it has been convincingly argued that the wine yeasts can originate from social wasps, where the symbiosis between wasps and yeasts is essential for the microbial transfer to wine ecosystems (Di Paola et al., 2023; Madden et al., 2018; Stefanini et al., 2012). Similarly, our results are reconcilable with the idea that bacteria for fermentation may originate from ants. This poses the question whether the ant holobiont was influential to the historic origins of yogurt or bread fermentation. Beyond traditional practices using ants in yogurt (Multu Sirakova, 2023; Öney Tan, 2010; Yalman, 1977), ant colony materials may be integrated in sourdough fermentation (Mutlu Sirakova 2023), providing further avenues for microbial transfer. Conceivably, the ant bacteria then could be backslopped in traditional fermentations, successively propagating and evolving in yogurts or sourdoughs. Although it is not clear when these traditions arose, it is noteworthy that the earliest evidence of yogurt and architectural depictions of ants and bees were found in prehistoric remains from our study region (Hendy et al., 2018; Walter, 2021). While these fragmentary anecdotes do not reconcile or resolve the origins of fermentative microbes, they highlight cultural threads that may connect ants and ferments.

The ant yogurt highlights the knowledge that can gained from traditional practices and ecological interactions, yet large-scale production of the yogurt itself may not be advisable. *Formica* red wood ants are of conservation concern due to recent declines in populations (Balzani et al., 2022). The bacteria within these ants, which can be isolated and used separately for fermentation, underscores the value of this biodiversity. However, public collection of ants is thus not sustainable. Furthermore, preparation of ant yogurt requires specialised knowledge. In modern gastronomy, *Formica* ants are typically frozen or placed in alcohol to kill a parasite the ants can carry, which is infectious to humans (Jensen et al., 2017). We found that fermentation started with frozen ants promote the growth of unfavourable bacterial communities, and that live ants must be used. In culinary applications with live ants, we then strained the starter with a microbiology-grade sieve to remove any potential parasites. In the traditional ant yogurt, knowledge holders may have maintained practices to prevent the spread of this parasite, but we did not address this in the scope of our research. Thus, careful consideration of both the cultural and biological aspects of traditional or historical practices is paramount when exploring their re-creation.

As a group of co-authors, including anthropologists, culinary innovators, and food scientists, we differentially perceive the implications of the ant holobiont on yogurt-making. From an anthropological viewpoint, it challenges the ethos of conventional yogurt fermentation. Unlike traditional yogurts, large-scale industrial yogurt production relies on a limited number of bacterial strains. Conversely, the lactic acid bacteria found in ant yogurt are maintained in the ants, yogurt, and environment, and are cultured through traditional dairy practices passed down over generations (Multu Sirakova, 2023). To preserve this microbial diversity, we must uphold its cultural heritage. In relation to modern gastronomy, elaborations on the use of ants in milk, along with their microbes and enzymes, may hold wider purchase for fermented foods. It opens new culinary possibilities for other dairy products and fermented beverages that are sour or effervescent. Beyond the possibility of imagining new foods, exposing the public to familiar foods made with microbes and insects could conceivably help change consumer perceptions of entomophagy and microbiology. From the food science perspective, microbes or enzymes from ant yogurts can be integrated into the food system to explore novel applications and flavours. For example, these ant microbes hold potential promise for new plant-based foods, such as dairy-free yogurts. While we differ in what the future of ant holobiont-facilitated fermentation may be, we agree it provides a powerful example that can broaden our thinking, as a society, about what is possible.

## Materials and Methods

### Traditional and modern culinary applications of ants

We travelled to Nova Mahala, Bulgaria to recreate ant yogurt with communities that retained memories and oral histories of the ant yogurt tradition. We based the yogurt fermentation on freshly acquired raw cow milk and *F. rufa* ants, from a colony found just outside of the village. The milk was warmed until nearly scalding, such that it could “bite your pinkie finger” (Multu Sirakova, 2023). Four live *F. rufa* ants were added to 400 mL of milk. A cheesecloth was secured over the milk, and fabric was wrapped around the glass container for insulation. The milk jar was then buried inside the ant colony, covering it completely with the mound material. The nest itself is known to produce heat (Coenen-Stass et al., 1980), and thus may act as an incubator for the yogurt fermentation. The milk was left within the colony and retrieved the next day, 26 hours later. The yogurt was then tasted, checked with pH strips, and stirred to observe coagulation.

Three culinary applications of the *Formica rufa* ants were developed by the research and development team at Restaurant Alchemist.

#### Ant yogurt ice cream sandwich

The “ant-wich” consisted of ant yogurt ice cream, ant gel filling, and ant tuile. To make the ant yogurt ice cream, organic 7% fat sheep’s milk (Hårbølle Mejeri, Møn, Denmark) was fermented with 1.6% live ants. Live ants were crushed and mixed in an aliquot of milk, which was passed through a strainer with pore size 100 µmM (Sigma Aldrich, Denmark) to remove ant body parts and potential parasites that may occur in low prevalence (Jensen et al., 2017). The strained aliquot was mixed with the rest of the milk, followed by an incubation period of 8 h at 40°C and subsequent storage at 4°C overnight. To sweeten and thicken the yogurt into ice cream, 300 g of sheep’s cream (Hårbølle Mejeri Møn, Denmark), and 22 g of sugar (Nordic Sugar, Denmark) was added to 380 g of ant yogurt, followed by 3 g of gelatine sheets (Condi, bloom 220, Denmark) and 5 g of ice cream stabilizer. The ant gel filling, based on an ant infusion, was made for the ice cream sandwich. For the infusion, dehydrated ants (*Formica rufa*) and water (filtered, pH 7) were mixed at the ratio of 10/100 (v/w), vacuum infused at 85°C for 8 h, strained through a fine mesh, and kept at 4°C. Subsequently, the ant gel was made with 100 g of ant infusion, 7.5 g of powdered glucose 33DE (Sosa, Spain), 70 g of inverted sugar (Trimoline, Cristalco, France), and 5 g of NH pectin (Sosa, Denmark). These were mixed, cooked for 5 min, and set to rest at 4°C for 24 h. Finally, an ant tuile was made to sandwich the ant ice cream and filling. The tuile was made by mixing 100 g of sweetened condensed milk (Tørsleffs, Denmark), 100 g of 00 wheat flour (Condi, Denmark), 100 g of egg whites, 3 g of baking powder (Condi, Denmark), 2 g of salt (Maldon), and 8 g of charcoal-black colorant (Sosa, Spain). The mixture was blended for 5 min, spread to measure 2 mm in height, and baked in the oven at 155 °C for 15 min. The final “ant-wich” was assembled to take the shape of an ant using a laser-cut stencil and served at -11 °C.

#### Ant mascarpone-like cheese

A mascarpone-like cheese was developed using ants as the coagulant. In common practice, lemon juice and milk or cream is added to make mascarpone, where the citric acid coagulates the dairy. Here, the acid from the ants likely contributed to the coagulation. Pasteurized sheep’s milk (7% fat) and cream (64% fat) (Hårbølle Mejeri, Møn, Denmark) were mixed with the proportions being 30% and 70%, respectively. The dairy mixture was homogenized and heated to 95°C. Then, 10 mL ant water infusion (described above) was added for each kilogram of this dairy mixture, stirring until complete coagulation (Munieweg et al., 2021). The curd was cooled to 4°C for 6 h and strained through a cheesecloth. The mascarpone- like cheese was stored at 4°C.

#### Cocktail clarified with ant milk-wash

Milk-wash is a technique commonly used in cocktails to clarify a liquid. Milk Punch, the common drink prepared with this technique, is a dairy-based drink dating to the late 1600s and early 1700s. The earliest written recipe of clarified milk punch comes from a 1711 cookbook by Mary Rockett (Wondrich, 2010). To do so, the milk is curdled by lowering the pH, which makes it possible to strain out the dairy solids. The result is a clear beverage with more richness and body. It is also a popular modern technique at cocktail bars, normally produced in large amounts (Wondrich, 2010). Using this technique, a cocktail was developed, but by inducing the milk coagulation with dehydrated ants. The formic acid replaced other products commonly used to lower the pH, such as lemon, lime, or any other acids like citric or ascorbic acid. For the cocktail, an alcoholic base was made using 181 g of brandy (Ximenez Spinola Battonage Brandy, Spain), 101 g of genepi liqueur (Genepi Grand Tetras, France), and 118 g of apricot liqueur (Bitter Truth Apricot Liqueur, Germany) mixed in a glass jar. 10 g of dehydrated ants (*Formica rufa*) were added and infused for 2 h at RT. The alcoholic mix plus dehydrated ants were poured into a glass jar with 200 g of milk (5% fat), stirred, and kept at 2 °C for 6 h to curdle the milk. Subsequently, the mixture was filtered through a coffee filter and kept at 2 °C until serving. The cocktail was garnished with 4 frozen ants. The cocktail is in the “aperitif” style of cocktails.

### Ant collection and processing for experimental yogurt

Experimental yogurts were made in spring, according to the tradition (Multu Sirakova, 2023), and in autumn, to test the seasonality of the fermentation. We used the red wood ant *Formica polyctena* for microbiome characterization and experimental yogurt fermentation, and *F. rufa* for microbiome characterization. *F. polyctena* is the sister species of *F. rufa*, where the later was used in the ethnographic account and at the Alchemist R&D. *F. polyctena* worker ants were collected at the end of May and the start of October 2022, from a colony in Rude Skov, Denmark (55.8334720, 12.474938410). Similarly, worker ants from a colony of *F. rufa*, found in Bøllemosen, Denmark (55.827676, 12.534667), were collected at the end of May and late September 2023. Ants were taken without disturbing the mound structure, considering several species within the group have experienced species declines within Europe (Balzani et al., 2022). Ants were gathered with sterilized equipment and kept on ice while transported to the lab.

*F. polyctena* worker ants were then processed to make the 3 elaborations of ant yogurt, live, frozen, and dehydrated. Live ants were provided sterile 15% sucrose as food and used to make ant yogurt within the week of collection. Prior to freezing, ants were aseptically removed from the nest material, then snap-frozen and stored at -20°C. Dehydrated ants were prepared by placing frozen ants in a vented Petri dish with sterile filter paper and incubated at 40°C for 16 h and were used within a week. Live and dehydrated ants were frozen at -20°C for later microbiome characterization.

The ant species were identified by barcoding of the *cytochrome c oxidase I gene (COXI)*. DNA was extracted with Chelex (Sigma Aldrich, Denmark), amplified, cleaned, and sequenced (Sinotte et al., 2021). PCRs were performed with primers LCO1490 5’- GGTCAACAAATCATAAAGATATTGG-3’and HCO2198 5’-TAAACTTCAGGRTGACCAAAAAATCA-3’ (Zuccon et al., 2012). PCR products were cleaned with PureIT ExoZap (Amplicon, Denmark) and sequenced at Macrogen (the Netherlands). Sequences were trimmed and aligned using default settings in Geneious (Biomatters Ltd., New Zealand), and the consensus sequence was compared to NCBI’s nucleotide database with BLASTn. The genetic identities of the consensus sequences were determined to be *F. polyctena* and *F. rufa*. This also matched the respective polydomous and monodomous nest architecture observed at the field sites and characteristic of the different species.

### Live, dehydrated, and frozen elaborations of experimental ant-yogurt

Ant yogurts were made under sterile conditions within a laminar flow hood, to assure that the ants were the sole source of microbes. The yogurt was made with 10% w/v reconstituted milk powder in distilled water (Sigma Aldrich, Denmark) and sterilized at 121 °C for 5 min to avoid caramelization. Yogurts were made in Falcon tubes with 30 mL of sterile milk and 0.5 g live ants, crushed with a sterile pestle, 0.5 g frozen ants, or 0.4 g dehydrated ants. Yogurts and negative controls were incubated at 42°C for 8 h and then stored at 4°C overnight, according to standard yogurt-making practices (Nagaoka, 2019; Robinson et al., 2006). The following day yogurts were subsampled for microbial culturomics, acid quantification, DNA extraction, and proteomics. Subsamples were stored at -20 °C.

In the spring, six samples of each of the live, frozen, and dehydrated ant-yogurts were made as well as 2 negative controls of milk alone, one incubated and one not incubated. In the autumn, the same design was used, but frozen ant-yogurt was not made as it was found to contain pathogens. Additionally, six samples of live and dehydrated at yogurt were made, where the milk with ants was passed through Whatman Filter Paper Grade 4 (Sigma Aldrich, Denmark), which retains particles up to 25 μm, and thus would theoretically remove any potential parasites while allow the bacteria to pass through. These filtered samples were only used for bacterial culturomics.

### DNA extraction of yogurt and ants

DNA was extracted from the yogurts and ants to assess the microbial community composition. All extractions were performed under sterile conditions in a laminar air flow. The yogurt samples were first homogenized by vortexing, then passed through a Corning cell strainer with pore size 100 µm (Sigma Aldrich, Denmark) to remove ant body parts. The yogurt underwent three washes prior to lysis to remove excess fats that may act as inhibitors and to pellet microbial cells. First, 3 g of yogurt and 10 mL of sterile 2% sodium citrate solution pH 7.4 were incubated at room temperature for 10 min, then centrifuged at 5,000 g for 10 min at 4°C. Second, the pellet was retained and washed in 15 mL TES buffer (50 mM Tris, 1 mM EDTA, 20% sucrose) and centrifuged at 5,000 g for 10 min at 4°C. A final wash was done in 1 mL TES, which was pelleted, and supernatant removed. For the ants, two ants were used per DNA extraction. Prior to extraction, 200 µL molecular water was added to the ants and they were crushed with a sterile pestle.

DNA extraction was performed with the DNeasy Blood and Tissue Kit (Qiagen, Germany) according to manufacturer’s instructions for gram positive bacteria with the following modifications. Physical lysis of ants and yogurts was performed via bead-beating with 0.1 mm G2 DNA/RNA Enhancer beads (VWR, Denmark) in a TissueLyser (Qiagen, Denmark) at 30 Hz for 3 min. Samples then underwent chemical lysis, including 30 min lysis in lysozyme with an addition of 2 µL of mutanolysin (25 U/μL) followed by 1 h proteinase K lysis. The final elution of the DNA extract was 200 µL in AE buffer, passed twice through the column to maximize yield. Due to exceptionally low DNA concentrations in some experimental yogurts, three extractions were performed per sample, and thus 9 g of yogurt in total was washed, extracted, and pooled per sample. Similarly, two ant extractions were pooled for a final sample representing four ants. The elutes were pooled and concentrated with AMP Pure XP Reagent (Beckman Coulter, Denmark) according to manufacturer’s instructions, then underwent a clean-up for inhibitors with the DNeasy PowerClean Pro kit (Qiagen, Denmark) according to manufacturer’s instructions.

The final samples for sequencing included yogurts, ants, and several controls. Yogurt samples thus included 20 samples from spring and 13 from autumn, as previously described. From the ants, there were six final samples for live and for dehydrated *F. polyctena* each from spring and autumn, and six samples of live *F. rufa* in spring and autumn. In addition to the experimental yogurts and ants, a mock community of standard bacteria (Zymobiomics, Nordic Biosite, Denmark) was included to determine any community bias generated by extraction and later metabarcoding. Further, three conventional plain yogurts were included as positive controls. Last, five blank DNA extractions were included as negative controls for DNA extraction.

### Quantification of bacterial load

The bacterial microbial loads of the yogurt, ant, and control samples were assessed with real-time PCR (qPCR). To quantify the bacteria in samples, the Femto Bacterial DNA Quantification Kit (Zymobiomics, Denmark) was used according to the manufacturer’s instructions. The Femto Bacterial Kit targets the 16S *rRNA* gene of bacteria and estimates the amount of bacterial DNA (ng/μL) in a sample based on a series of standards and non-template control. qPCR of samples was run in duplicate on Mx3005p (Agilent, Germany). The technical replicates demonstrated minimal variation and were thus averaged. Subsequently, the amount of DNA/mL of yogurt, in the total yogurt, or ant starter, was back calculated based on the standards and experimental parameters.

### Bacterial community metabarcoding and analysis

The bacterial community of the ants, yogurts, and controls was characterized with bacterial metabarcoding. Library preparation and targeted sequencing of the 16S *rRNA* V3-V4 region was performed with the primers 341F (5’ -CCTAYGGGRBGCASCAG- 3’) and 806R (5’ - GGACTACNNGGGTATCTAAT- 3’) at Novogene Europe (Cambridge, United Kingdom). In brief, amplicons for the targeted region underwent size selection, then equivalent amounts of PCR product were end-repaired, A-tailed, and ligated with Illumina adapters. Library concentrations were checked with Qubit and qPCR, and size distribution was checked with BioAnalyzer. Libraries were pooled and sequenced on Illumina for pair-ended 250 bp at a depth of 30k per sample. After sequencing, primers and adaptors were removed by Novogene, and raw reads were provided for analysis.

The analysis of sequences was carried out in R (v. 4.3.1) (Team, 2023), which included determination of amplicon sequence variants (ASVs), taxonomic assignment, visualization, and statistical analyses. The dada2 pipeline (v. 1.30.0) (Callahan et al., 2016) was used for ASV and taxonomic assignment. Reads were not trimmed due to the overall high quality of sequences, but were filtered with the default parameters, with *maxEE* adjusted to (2,4). Reads were then paired with *mergePairs*, *minOverlap* set to 16, and chimeras were removed. Taxonomy was assigned to the species level where possible with the SILVA database (v. 138.1) (Glöckner et al., 2017). Potential contaminants were then removed using SCRuB (v.0.0.1) (Austin et al., 2023) based on the blank negative controls that were successfully sequenced. Mitochondria, chloroplasts, and Eukarya were removed, resulting in 63.87% of ASVs with genus-level classification and 82.23% with family-level. The mock community was then assessed to determine any potential bias created by DNA extraction and sequencing. All eight bacterial genera present in the mock community were validated.

The metabarcoding data was then analysed and visualized using the packages phyloseq (v.1.46.0) (McMurdie & Holmes, 2013), microviz (v.0.12.0) (Barnett et al., 2021), and ggplot2 (v.3.4.4) (Wickham, 2011). The composition of the microbiome of ants and yogurts was visualized with microviz. The beta diversity of the yogurts was visualized with phyloseq through a NMDS ordination of their Bray-Curtis distances, resulting in a plot ordinated by yogurt treatments. The alpha diversity of the yogurts was further determined with phyloseq. Last, the microbial loads of Lactobacillaceae and Bacillaceae in the ant starters and yogurts were estimated by multiplying the relative abundance of each genus, based on the metabarcoding data, by the quantified bacterial load, based on 16S qPCR. This was visualized using ggplot2.

We assessed the effect of treatment (live, dehydrated, or frozen) on the beta diversity of the yogurts using Bray-Curtis distances and an adonis PERMANOVA using the package vegan (v.2.6.4) (Oksanen et al., 2013) and function *adonis2*. Subsequently, we performed a pairwise-comparison between the three treatments based on the PERMANOVA using the package pairwiseAdonis (v.0.4.1) (Martinez, 2017), which performs *p*-value adjustment for multiple comparisons. The alpha diversity of the three yogurts was based on the Shannon index. We performed a general linear model to determine the effect of treatment on the alpha diversity, confirming the parametric assumptions of the model. Pairwise comparisons were then performed with a Tukey HSD. Pairwise comparisons of the Lactobacillaceae and Bacillaceae load in the ant starters and yogurts was performed with two sample t-tests for equal or unequal variance depending upon the data.

### Bacterial culturomics of milk ferments and ants

Bacteria from the ant yogurts and ants were cultured and identified with Sanger sequencing. 100 µL of yogurt sample was inoculated and plated in De Man, Rogosa and Sharpe (MRS) (VWR Chemicals, US), Gifu Anaerobic Medium, modified (GAM) (HyServe GmbH & Co. KG, Germany), yeast peptone dextrose (YPD), and Luria-Bertani (LB) media. YPD contained 10 g/L yeast extract, 20 g/L peptone, and 20 g/L D-glucose. LB contained 10 g/L peptone, 5 g/L yeast extract and 0.5 g/L sodium chloride. A total of 20 g/L agar was added for solid cultivations. For ants, individual ants were inoculated into liquid media. Tubes and plates were incubated at 30°Cover 2 days, or until growth was visible. All four media were cultivated in aerobic conditions and MRS and GAM were also incubated anaerobically. Subcultures were plated and incubated in the same conditions. Distinct bacterial colonies from plates were identified with Sanger sequencing of the conserved 16S *rRNA* gene, which included colony PCR (cPCR) with primers 5’- AGAGTTTGATCCTGGCTCAG-3’ and 5’-CCGTCAATTCCTTTRAGTTT-3’ (Cebeci & Gürakan, 2008). All cPCR reactions consisted of 12.5 µL of Red-Taq Mastermix (VWR, Denmark), 0.5 µL of forward 10 µM primer, 0.5 µL of reverse 10 µM primer, 11.5 µL of water and a bacterial colony. The reactions followed the same thermocycler program that consisted of an initial 4 min at 94 °C, followed by 35 cycles of 94 °C for 30 s, 51 °C for 30 s, 72 °C for 1 min with a final 72 °C for 10 min. PCR products were Sanger sequenced (Eurofins Scientific SE). Sequences were trimmed and aligned using default settings in Geneious (Biomatters Ltd., New Zealand), and the consensus sequence was compared to NCBI’s nucleotide database with BLASTn. Results were presented in a Sankey diagram (app.rawgraphs.io).

### Quantifying pH, lactic, acetic, and formic acid in yogurt

The pH and coagulation of all yogurts were assessed before and after fermentation. After fermentation, the pH was measured from one sample of each of the groups, and the coagulation of the treatment group was assessed. Given the abundance of formic acid found in *Formica* ants (O’Rourke, 1950), we also examined the effect of formic acid on coagulation. Formic acid (Sigma Aldrich, Denmark) was added to the sterile milk until pH 4.6, 5.15, 5.9, and 6.6, which represented the range of pHs observed in conventional yogurts, ant yogurts, and sterile milk. The pH of the negative control was also measured. Additionally, pH of all the samples was measured prior to DNA extraction and enzyme analysis, after they had undergone a cycle of freezing and thawing. This was performed on a separate pH reader than that which provided the initial measurement before and after fermentation.

The amount of formic, lactic, and acetic acid in the samples was quantified with high-performance liquid chromatography (HPLC). Yogurt samples were centrifuged at 5000 rpm for 5 min at 4°C, and their supernatant was passed through a 0.2 μm syringe filter (Sigma Aldrich, Denmark). The filtrate was analysed by a Dionex Ultimate 3000 HPLC system (Thermo Fisher Scientific, USA) equipped with a refractive index detector and a Bio-Rad Aminex HPx87 column (Bio-Rad, USA). The column oven temperature was set at 30°C. The flow rate was set at 0.6 mL/min with 5 mM H₂SO₄ (Sigma Aldrich, Denmark) as an eluent. The injection volume was 10 μL. Peaks were identified by comparison to the prepared standards, and integration of the peak areas was used to quantify acids from obtained standard curves using the software Chromeleon7 (Thermo Fisher Scientific, USA). Plots were visualised in GraphPad Prism. T-tests were used to compare organic acid levels between samples, with Benjamini-Hochberg correction for multiple testing.

### Proteomics of ant yogurts

Sample preparation for proteomics analysis was performed directly from the supernatant of the yogurt samples after centrifugation for 10 min at 14,000 g and further processed as described earlier (Kozaeva et al., 2021). After centrifugation, the supernatants were subjected to the bicinchoninic acid (BCA) assay to estimate protein concentrations. Trypsin and LysC digestion mix (Promega) was added to 20 μg protein of each sample and incubated for 8 h. Trifluoroacetic acid was added to halt digestion and the samples were desalted using C18 resin (Empore, 3M) before HPLC-MS analysis.

HPLC-MS analysis of the samples was performed on an Orbitrap Exploris 480 instrument (Thermo Fisher Scientific) preceded by an EASY-nLC 1200 HPLC system (Thermo Fisher Scientific). For each sample, 1 μg of peptides was captured on a 2 cm C18 trap column (Thermo Fisher 164946). Subsequently separation was executed using a 70 min gradient from 8% (v/v) to 48% (v/v) of acetonitrile in 0.1% (v/v) formic acid on a 15-cm C18 reverse-phase analytical column (Thermo EasySpray ES904) at a flow rate of 250 nL/min. For data-independent acquisition, the mass spectrometer was run with the HRMS1 method (Xuan et al., 2020) preceded by the FAIMS Pro Interface (Thermo Fisher Scientific) with a compensation voltage (CV) of -45 V. Full MS1 spectra were collected at a resolution of 120,000 and scan range of 400-1,000 m/z, with the maximum injection time set to auto. MS2 spectra were obtained at a resolution of 60,000, with the maximum injection time set to auto and the collision energy set to 32. Each cycle consisted of three DIA experiments each covering a range of 200 m/z with a window size of 6 m/z and a 1 m/z overlap, while a full MS scan was obtained in between experiments.

To assign the proteases to their potential organisms of origin, we created a database consisting of the proteases, peptidases and proteins based on UniProt reference proteomes, the gold standard in proteomics. The proteomes consisted of the ant host, bacteria from the ant yogurt, and two conventional yogurt bacteria. For the ant, we used the *F. execta* proteome, since the *F. polyctena* proteome was not publicly available, and *F. execta* is a closely related species. From the *F. execta* proteome, we selected the proteases specifically. For all the bacteria included, we refined our databased to only include proteases and peptidases in the proteome. We selected bacteria most prevalent among the bacteria isolated from the live ant yogurt bacterial metabarcoding and culturomics: *Fructilactobacillus sanfranciscensis*, *Oecophyllibacter saccharovorans*, and *Paenibacillus sp.* We also included *Wolbachia pipentis*, which is closely related to the obligate intracellular bacteria found in *Formica* ants (Jackson et al., 2022) and is represented in our analysis. We included the conventional yogurt bacteria *Streptococcus thermophilus* and *Lactobacillus delbrueckii* subsp. *bulgaricus. Streptococcus* was only present in low abundance in the bacteria metabarcoding data, and *L. delbrueckii* subsp. *bulgaricus* was isolated in one instance. We included them to determine if the ant yogurt contained proteases with the same or similar functionality. Last, we included common milk proteins, such as caseins, found in the *Bos taurus* proteome. For DIA data analysis, Spectronaut v18 (Biognosys, Switzerland) was used for protein identification and relative quantification of peptides. The default settings for “directDIA” were applied with an FDR cut-off of 1%, except for MS1 quantification for the peptides. Protein abundances were inferred from the peptide abundances using the MaxLFQ algorithm available within Spectronaut.

In the analysis of the results, first we determined if proteases, peptides, and milk proteins, changed in relative abundance between spring or autumn and live or dehydrated samples. We were unable to determine the abundance of a subset of proteases identified, likely due to their low abundance, and they were therefore not included in the analyses. For each protein, the average relative abundance was computed across all autumn samples and all spring samples. Similarly, the average relative abundance was computed across all live samples and all dehydrated samples.

The log2 fold change (log2FC) was calculated using the formula: log2FC=log⁡2(average abundance in condition 1/ average abundance in condition 2) *P*-values were obtained by comparing the relative abundances of each protein across all samples and their replicates in the autumn versus spring conditions. Similarly, *p*-values were calculated by comparing the relative abundances of each protein across all samples and their replicates in the alive versus dehydrated conditions. The Benjamini-Hochberg procedure was applied to adjust the *p*-values for multiple comparisons, controlling the false discovery rate (FDR). The adjusted *p*- values were then transformed to their -log10 values for further analysis and visualization.

Second, we visualized the composition of the proteases and peptidases in GraphPad Prism. To do so, we removed the milk proteins originating from *Bos taurus*, given they made up a high proportion of the proteome. The aim was thus to clarify ant or bacterial contributions to the enzymatic profile. The relative abundance of each protease or peptidase visualized was thus based on the total abundance of proteases and peptidases in each sample, and not the proteome.

## Data availability

Amplicon sequences from metabarcoding of ants and yogurts are available on SRA (BioProject PRJNA1162421) and Sanger sequences of isolated bacterial strains (PQ535580-PQ535692 and PQ651799-PQ651879) and ants (PQ453570-PQ453574) are available on Genbank. Proteome data will be made available on the ProteomeXchange database under identifier identifier PXD057843. Additional data frames have been deposited on FigShare and will be made public prior to publication (https://figshare.com/s/a570abbd84d0fb09aa83).

## Code availability

The code for analyses is available on GitHub and will be published on Zenodo once accepted (https://github.com/vsin632/ant_yogurt.git).

## Author contributions

Conceptualization: V.M.S., L.J.J., R.R.D., V.R-V., D.P.V., S.M.S; Methodology: V.M.S., V.R.-V., D.P.V, N.R.V, S.K.T.P, J.G., E.M.V, S.B.A; Formal analysis: V.M.S., V.R.-V., S.K.T.P; Investigation: V.M.S., V.R.-V. D.P.V., S.M.S., N.R.V., A.C-M., S.K.T.P., J.G., E.M.V., D.Z.; Resources: L.J.J., S.B.A, R.M., S.M.S.; Data curation: V.M.S.; Writing – original draft preparation: V.M.S., V.R-V., L.J.J., R.R.D; Writing – review & editing: all authors; Visualisation: V.M.S., V.R.- V.; Supervision: L.J.J., R.D.D., S.B.A., R.M.; Project administration: V.M.S., L.J.J.; Funding acquisition: S.B.A., L.J.J., R.R.D., R.M.

## Acknowledgments

We thank Rasmus Stenbak Larsen for the assistance in collecting the ant colonies. We acknowledge myrmecologist Dr. Vera Anatova from the Bulgaria Academy of Sciences, who provided knowledge and assisted us with the permit НС ЗП-122/18.04.2023 from the Ministry of Environment and Water to collect wood ants in Bulgaria. We thank villagers from Nova Mahala for sharing their knowledge, especially Musserem, Mehmet, and Fatma. We acknowledge gastronomers Fejsal Demiraj and Tangör Tan for their personal communications around the traditional use of ants in dairy fermentations. Last, we thank the laboratory staff at the Globe Institute for their expert support and guidance, especially Lasse Vinner, Tina Blumensaadt Brand, and Pernille Selmer Olsen. V.M.S., A.C.M., J.G., and S.B.A. were supported by the Danish National Research Foundation Centre for Evolutionary Hologenomics DNRF 143. V.R.V, and L.J.J. were supported by the Novo Nordisk Foundation, NNF Grant number: NNF20CC0035580.

