## Supplemental Figures for "Making yogurt with the ant holobiont uncovers bacteria, acids, and enzymes for food fermentation"

**Supplemental Information**

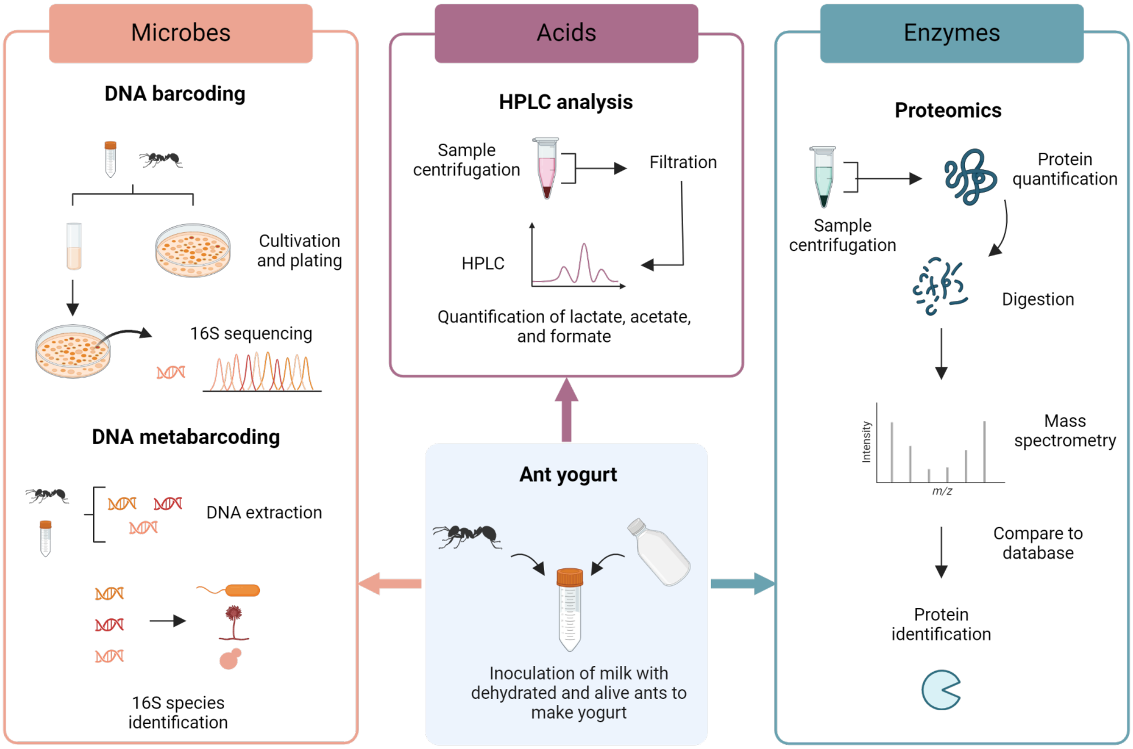


**Figure S1. Experimental workflow to characterize microbes, acids, and proteases in ant yogurt.** Milk was inoculated with live, frozen, or dehydrated ants and incubated overnight. Microbes present in the yogurt and ants were identified with culturing and DNA barcoding as well as metabarcoding. The amounts of the acids lactate, acetate, and formate in the yogurts as well as controls were quantified by HPLC. The presence and prevalence of proteases and peptidases in the yogurts originating from ants and microbes was characterized with proteomics. (The figure was created with BioRender.com).

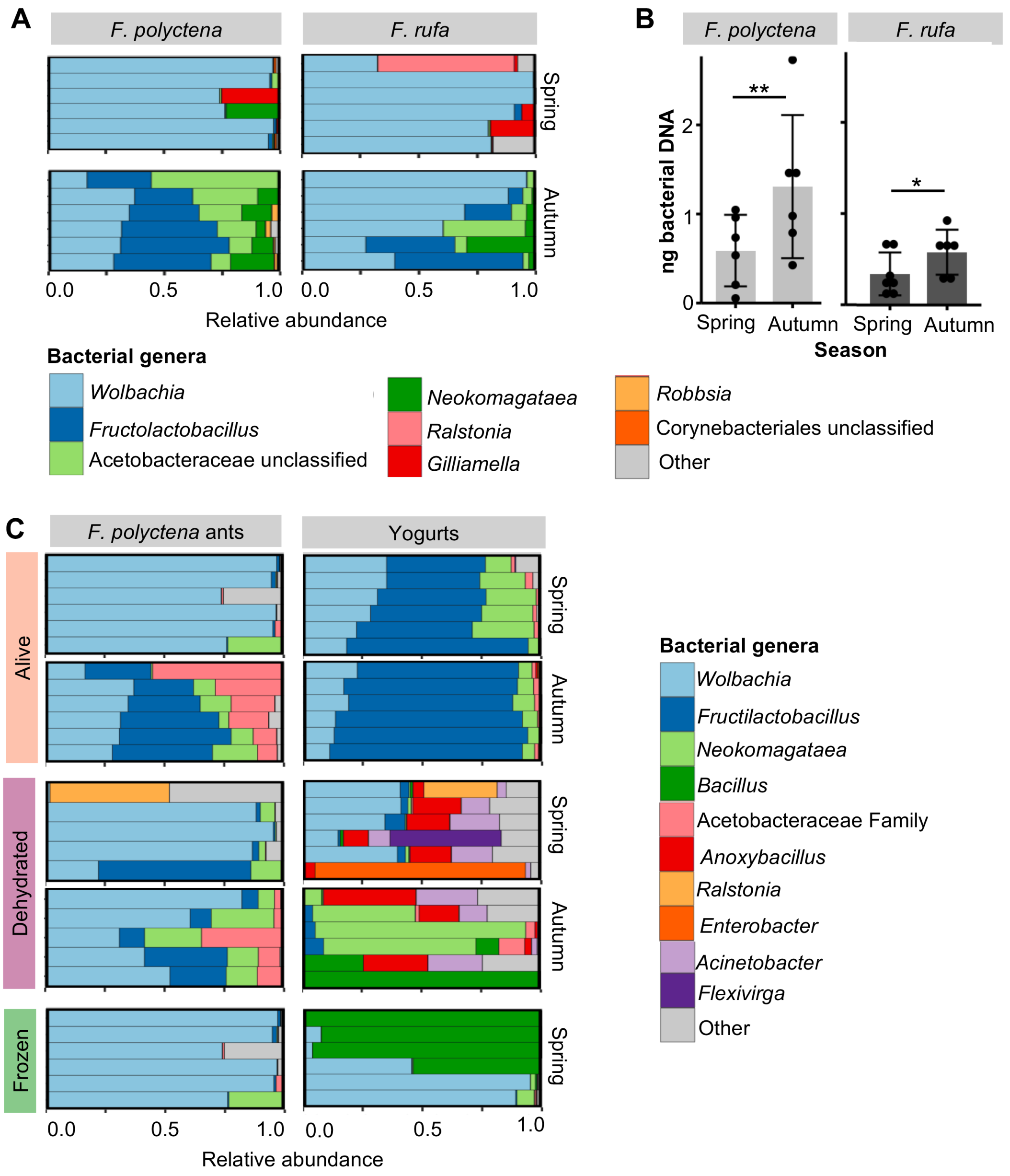


**Figure S2. Genus-level composition of microbiomes and seasonal bacterial load of ants.** (**A**) The genus-level microbiome composition of both ant species across seasons, where each bar represents a replicate of four pooled ants from a colony of each respective species. (**B**) The absolute abundance of bacteria biomass in *Formica* ant species across seasons, where each dot represents a replicate. Asterisks indicate significant differences based on pairwise t-tests (* = $p \leq .05$; ** = $p \leq .01$) (**C**) The genus-level bacterial microbiome of ants across the three preparations (live, dehydrated, and frozen) and corresponding ant yogurts made in late spring and early autumn. The bars represent replicates. Note that the live ant microbiomes are also represented in panel **(A)**, and similarly the live and frozen ants are identical, given all samples were frozen prior to analysis. They are shown again for direct comparison to the corresponding yogurt microbiomes.

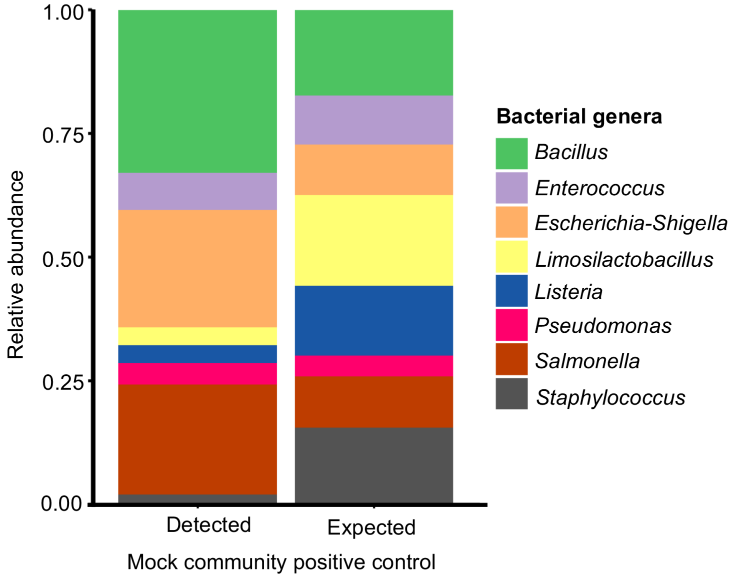


**Figure S3. Detection bias of bacterial abundances based on mock community control.** Mock community standards are commonly included as positive controls in microbiome studies to assess bias caused by DNA extraction and primer specificity. The detected relative abundances compared to the expected mock community standard abundances indicates that bacteria related to *Bacillus*, *Escherichia-Shigella*, and *Salmonella*, may be overestimated within the metabarcoding data compared to their true abundance. Similarly, *Enterococcus*, *Limosilactobacillus*, and *Listeria* may be underestimated within the metabarcoding data compared to their true abundance.

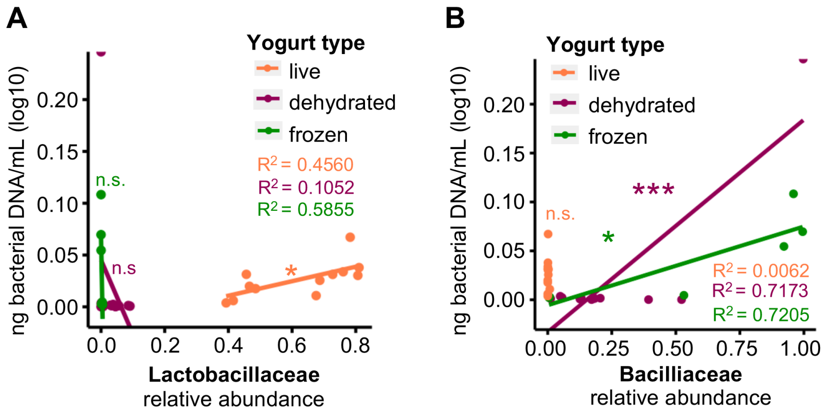


**Figure S4. Bacterial load and relative abundance of Lactobacillaceae and Bacillaceae.** (**A**) The relative abundance of lactic acid bacteria (Lactobacillaceae) and the total bacterial biomass for each sample, indicating little to no lactic acid bacteria was found in dehydrated and frozen ant yogurts. (**B**) The relative abundance of Bacillaceae and the total bacterial biomass for each sample, indicating little to no Bacillaceae was detected in live ant yogurts. Dots represent individual samples. Lines, $R^2$, and *p*-values are based on linear models for each yogurt type (* = $p \leq .05$; ** = $p \leq .01$; *** = $p \leq .001$; n.s. = p>0.05).

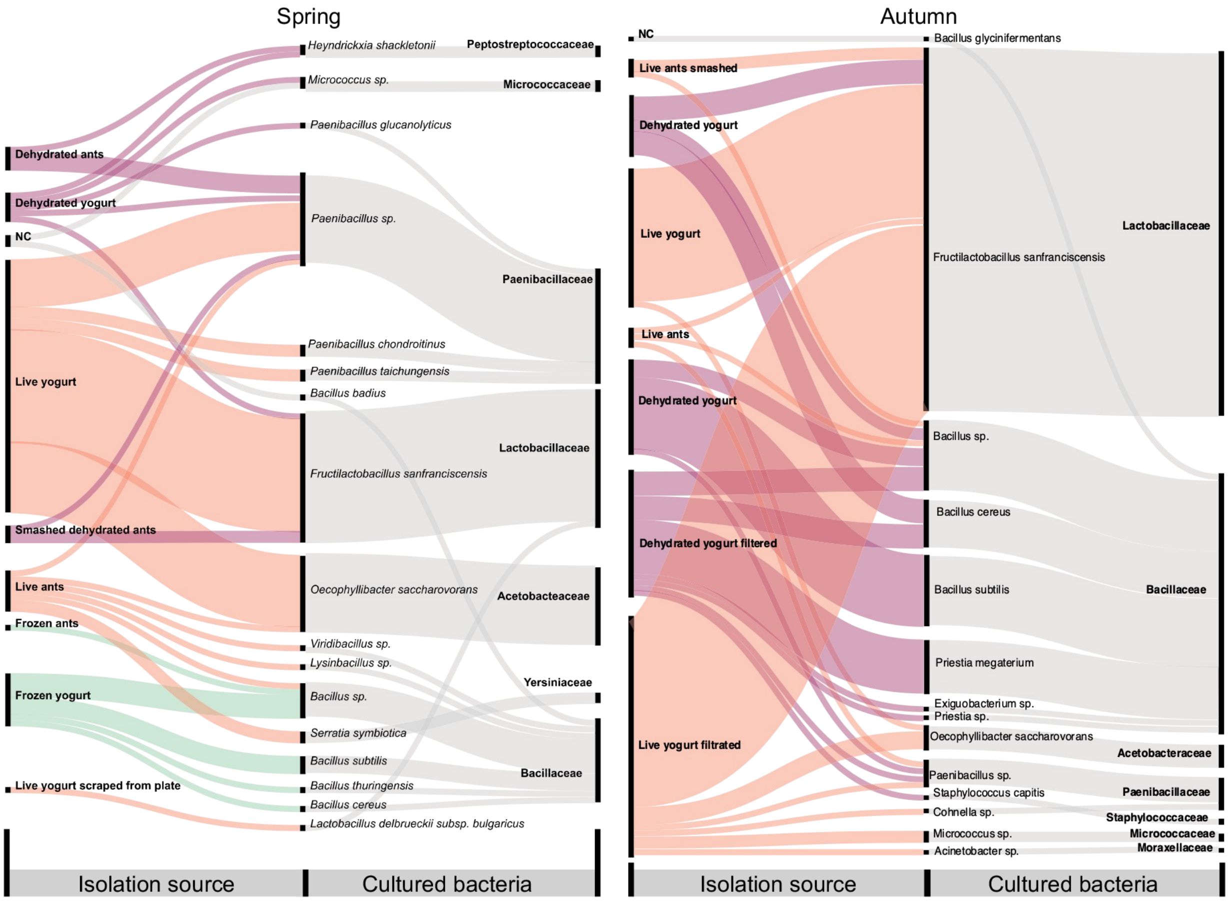


**Figure S5. All bacteria cultured from ants and ant yogurts in spring and autumn.** Three biological replicates of ant yogurts and ants were grown in general LB and YPD media aerobically, and selective MRS and GAM media aerobically and anaerobically. They were directly plated from the yogurt and grown in liquid media with subsequent plating. Bacterial colonies were sampled, 16S *rRNA* maker genes sequenced, and taxonomically identified. The thickness of each line represents the number of plates where the bacteria grew compared to other samples.

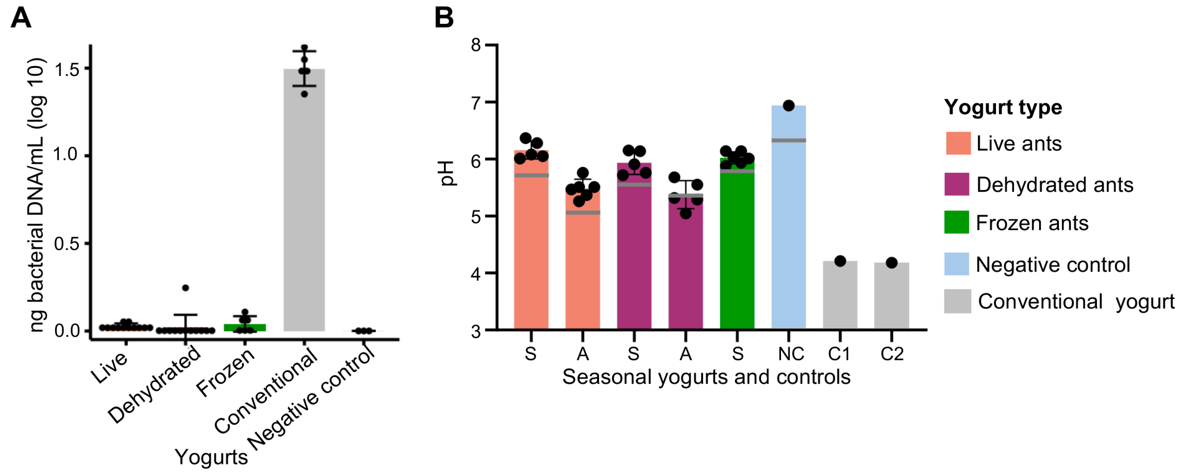


**Figure S6. Bacterial load and pH of ant yogurts, conventional yogurts, and controls.** (**A**) The bacterial load, as detected by qPCR targeting the 16S *rRNA* gene. Error bars indicate standard error. (**B**) pH of yogurts across yogurt types. Seasonal yogurts from spring (S) and autumn (A), the negative control of milk alone, and two conventional yogurts were measured. The grey line indicates initial measurement of the pH of one yogurt immediately after the fermentation, and the dots indicate later pH measurement of all yogurts after a freezing and thaw cycle. The measurements indicate consistent trends, but initial measurements are slightly lower potentially because they were conducted on a different pH meter or prior to freeze-thaw of the yogurts. Error bars indicate standard error.

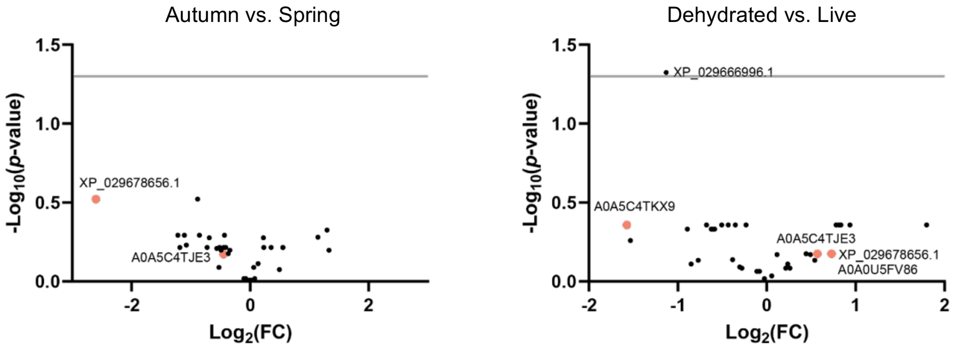


**Figure S7. Differential abundance of ant yogurt proteases and peptidases across seasons and elaborations.** Fold change (FC) of autumn compared to spring and dehydrated compared to live ant yogurts plotted against the Benjamini Hochberg corrected *p*-value. The grey line represents the threshold of significance (α=0.05). In orange are highlighted proteases with documented evidence of cleaving casein proteins.

**Table S1.** pH of yogurts and formic acid controls before and after incubation.

**Table S2.** Absolute abundance of milk proteins, proteases, and peptidases in proteomes.
